# RAPID: Reliable and efficient Automatic generation of submission rePorting checklists with Large language moDels

**DOI:** 10.1101/2025.02.13.638015

**Authors:** Zeming Li, Xufei Luo, Zhenhua Yang, Huayu Zhang, Bingyi Wang, Long Ge, Zhaoxiang Bian, James Zou, Yaolong Chen, Lu Zhang, the ADVANCED working group

## Abstract

**Importance:** Medical reporting guidelines are significant in improving the transparency, quality, and integrity of medical research, particularly in randomized clinical trials; adherence to these guidelines supports research interpretability and has direct implications for downstream applications, such as patient treatment. However, with over 600 distinct reporting guidelines, manual assessments are often time-consuming and labor-intensive.

**Objective:** To evaluate an automated reporting checklist generation tool using large language models and retrieval augmentation generation technology, called RAPID.

**Design, Setting, and Participants:** This study used large language models to design a retrieval augmentation generation architecture and collected published journal articles as training and validation sets to optimize prompts within the framework and comprehensively evaluate the performance of the framework. Medical reporting experiments were collected from 50 randomized controlled trials without the intervention of AI tools and 41 randomized controlled trials with the intervention of AI tools.

**Main Outcomes and Measures:** For effective evaluation of the performance of this tool, a classification accuracy metric (Reported/Not Reported) defined as the number of correct judgments divided by all judgments and a content consistent score metric defined as the number of contents retrieved by the tool that are the same as those retrieved by researchers divided by the total number of judgments were calculated.

**Results:** The RAPID tool uses the widely used Word document and Portable Document Format as an input file. Fifty published journal articles without the intervention of AI tools and 41 published journal articles with the intervention of AI tools were collected as CONSORT and CONSORT-AI datasets. All of the CONSORT reporting items (37) were included in the tool. RAPID achieved a high average accuracy rate of 92.11% and a content consistency score of 81.14% on the CONSORT dataset. Of the CONSORT-AI reporting items, 11 items related to the intervention of AI tools were included in the tool. RAPID achieved an average accuracy of 83.81% with a content consistency score of 72.51% on the CONSORT-AI dataset. For these two reporting guidelines, a training set of 5 articles was selected from each dataset to refine the prompts used in the tools for CONSORT and CONSORT-AI reporting checklist. The validation set of the remaining articles was used to assess the performance of the RAPID. The RAPID tool used the Word document and Portable Document Format of the articles as input files. A RAPID graphical user interface was built using JavaScript and Vue.

**Conclusions and Relevance:** The RAPID tool is designed to assist in the reporting of various types of trials. RAPID has strong scalability, which can be easily adapted to different medical reporting guidelines without transfer learning on a large dataset. RAPID may effectively save time and improve working efficiency for different user groups, for example, 1) simplifying the submission process and improving report quality by verifying manuscript completeness for medical authors; 2) facilitating evaluation of report quality for medical researchers; 3) expediting manuscript distribution for medical editors; and 4) identifying reporting deficiencies and providing deeper insights for review comments for reviewers.

**Key Points:** *Question:* Can large language model tools automatically generate medical reporting checklists for manuscripts of different types of clinic trials?

*Findings:* An automated reporting checklist generation tool using large language models and retrieval augmentation generation technology, RAPID, was developed. RAPID was fully evaluated on all items across two separate datasets related to the Consolidated Standards of Reporting Trials (CONSORT) and CONSORT-AI reporting guidelines. In the first dataset corresponding to CONSORT, RAPID achieved an average accuracy of 92.11%, while in the second dataset associated with CONSORT - AI, it reached an average accuracy of 83.81%. Additionally, RAPID is highly scalable. It can be easily and smoothly adapted to different medical reporting guidelines without transfer learning on a large dataset.

*Meaning:* RAPID can effectively save time and improve working efficiency for different user groups such as medical authors, researchers, editors, and reviewers.

## 1 Introduction

Medical reporting guidelines, such as CONSORT (**Supplementary eTable 1**), are critical standards for writing and publishing medical research reports, which are required to cover the content corresponding to each item in the guidelines. Transparent and standardized reporting improves the interpretability of medical studies, particularly randomized clinical trials (RCTs), which form the backbone of evidence-based clinical care and drive future research. Accurate, comprehensive, and scientifically sound reporting can uphold the integrity and transparency of different types of downstream studies such as the creation of systematic reviews, which in turn improves the overall quality of medical evidence and ultimately contributes to better patient outcomes ^18^.

The Enhancing the QUAlity and Transparency Of health Research (Equator) Network^2^ currently hosts over 600 medical reporting guidelines. With such an extensive range of standards, it is unrealistic for researchers to be familiar with all of them. In addition, manually reviewing medical literature for compliance with these guidelines is both time-consuming and labor-intensive. For example, evaluating an observational study against the 22 items of the STROBE (STrengthening the Reporting of OBservational studies in Epidemiology) checklist takes a median of 30 minutes (range: 15–40 minutes) ^15^. Furthermore, new extensions to these guidelines are frequently proposed ^11^, aggravating the challenge ^25^. These barriers highlight the need for automated tools to assist researchers in efficiently checking medical reporting items, improving both the speed and quality of research reporting.

Recent advancements in artificial intelligence offer promising solutions to these challenges. While traditional Natural Language Processing (NLP) ^5,19^ has been the primary technology used for tasks like information retrieval and summarization, more recent developments in Generative Pre-trained Transformer-based Large Language Models (GPT-based LLMs) ^17^ and Retrieval-Augmented Generation (RAG) ^3,9,10,13^ have significantly improved the efficiency and accuracy of these tasks. GPT-based LLMs, leveraging the Transformer architecture ^26^, are pre-trained on vast amounts of text and fine-tuned using reinforcement learning based on human feedback ^21^. This process empowers these models to excel in complex semantic comprehension tasks and makes them highly suitable for nuanced text analysis in medical reporting.

In 2020, Wang et al. developed CONSORT-NLP ^27^, an automated tool designed to generate CONSORT reports and assess adherence to CONSORT guidelines with an impressive accuracy of 90%, completing assessments in under 30 seconds per paper. However, the tool faced several limitations: (1) It only considered 34 out of the 37 items in CONSORT as the remaining three were too complex (complex items) for the method to answer accurately; (2) It employed traditional NLP models which operated as a ‘black box’, lacking transparency, so users could not understand the reasoning behind decisions; and (3) The tool lacked flexibility, requiring retraining on specific datasets for different reporting guidelines, which limited its wide applicability.

In this study, we introduced RAPID, a novel framework designed to overcome these limitations by integrating open-source GPT-based LLMs and RAG. RAPID leveraged the robust semantic capabilities of LLMs to break down checklist items into sub-queries for more accurate information extraction and selection of relevant paragraphs. Moreover, the natural language understanding capability of LLMs allowed for the generation of human-like explanations of the results, enhancing transparency and increasing user trust in the method. Furthermore, our proposed multi-agent architecture ensured the framework’s adaptability to a wide range of medical reporting guidelines with minimal performance loss, without the need for retraining on specific datasets.

We manually annotated 50 and 41 RCT research papers based on CONSORT and CONSORT-AI checklists ^16,22^ (**Methods 2.1**), respectively, and regarded them as evaluation datasets (CONSORT and CONSORT-AI dataset). After a comprehensive comparison with the Vanilla Prompt Method (**Methods 2.3**), which utilized multiple popular closed-source LLMs, our framework implemented with an open-source LLM, deepseek-chat ^6^, showed state-of-the-art performance on both datasets. We achieved an accuracy of 92.11% with a content consistency score (CCS) (**Methods 2.5.2**) of 81.14% on the CONSORT dataset, and an accuracy of 83.81% with a CCS of 72.51% on the CONSORT-AI dataset. However, the Vanilla Prompt Method combined with ChatGPT-4o ^1^ only achieved an accuracy of 84.59% with a CCS of 77.51% on the CONSORT dataset, and an accuracy of 73.84% with a CCS of 62.75% on the CONSORT-AI dataset.

These findings highlighted not only the robustness and scalability of our framework but also demonstrated the advantage of leveraging open-source models for more efficient and effective analysis. Additionally, we have developed a user-friendly web-based interface that improved transparency by providing users with clear reasoning behind each decision, thereby increasing interpretability and user confidence. The latest system version has been deployed at http://rapid.advancedguides.ai.

## 2 Methods

### 2.1 RCT Paper Selection and Gold Standard Extraction

For this study, we developed two datasets, each aligned with a specific case study:

(1) **Dataset for CONSORT checklist:** This dataset comprises 50 randomized controlled trials (RCTs) randomly selected from PubMed, all published within the past five years; and (2) **Dataset for CONSORT-AI checklist:** This dataset consists of 41 RCTs involving AI interventions, drawn from a previously published systematic review^16^.

We manually evaluated the papers in the CONSORT dataset against all items in the CONSORT checklist. For the CONSORT-AI dataset, we assessed the papers against only the 11 AI-related items from the CONSORT-AI checklist. Pairs of researchers verified whether each item in the articles was ‘Reported’ or ‘Not Reported’ and extracted the corresponding paragraphs based on the checklist descriptions to support classification.

### 2.2 Prompt Engineering

Recent advances in GPT-based large language models have demonstrated strong performance in complex reasoning tasks, with prompt engineering being a crucial component. Specifically, Chain of Thought (CoT) prompting^28,30^ has emerged as a powerful technique, categorized into two main approaches: (1) **Zero-shot CoT Prompt:** This encourages LLMs to generate intermediate reasoning steps by using a simple instruction, such as “Let’s think step by step”; and (2) **Few-shot CoT prompt:** This provides LLMs with explicit examples of intermediate reasoning steps to guide the problem-solving process. In general, effective prompts, containing detailed instructions and contextual domain information, significantly enhance the LLM’s understanding of complex tasks in specific fields. In this study, we designed our prompts based on ChatGPT-3.5, which included role definitions, task descriptions, inference steps, response formats, and illustrative examples (**Supplementary eFigures 5-8**). We iteratively refined these prompts through trials on five randomly selected RCTs, adjusting them until the outputs were satisfactory. The finalized prompts were then used in the main experiments.

### 2.3 Vanilla Prompt Method

The Vanilla Prompt Method used pre-constructed prompt templates to input checklist items and papers into the LLMs. These prompt templates guided the LLMs to extract relevant information and classify items as either ‘Reported’ or ‘Not Reported’ within the literature.

### 2.4 Article Preprocessing

In the experimentation, to prepare the selected RCTs for analysis, original PDF documents were first converted into editable Word format using the online tool Smallpdf (https://smallpdf.com). Next, tables and flowcharts were converted to LaTeX format using the OpenAI ChatGPT API. In addition, irrelevant elements, such as author names, institutions, and references, were removed from each article to reduce token usage during experiments. Following this, the articles were segmented into individual sentences, and each sentence was transformed into an embedding using a semantic model, such as bge-large-en^29^. After that, adjacent sentences close in the embedding space were merged to form paragraphs, with the process ensuring semantic consistency and avoiding redundant information within each paragraph.

### 2.5 Statistical metrics

#### 2.5.1 Accuracy

As detailed earlier (**Methods 2.1**), each RCT was manually evaluated against the corresponding reporting guideline. The gold standard for each item was categorized as either ‘Reported’ or ‘Not Reported,’ and the corresponding original content was extracted from the literature. To evaluate the performance of judgment consistency, we calculated accuracy, referred to as the Overall Consistency Score in the field of medical reporting. Accuracy is calculated as follows:

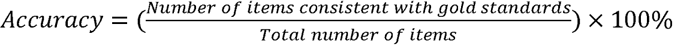

#### 2.5.2 Content Consistency Score

In real-world scenarios, different human researchers conducting checklist evaluations may consistently judge certain items as "Reported," but the specific passages they refer to may vary. Therefore, accuracy alone may introduce bias, as it can not reflect how consistently the content produced by models aligns with information extracted by human researchers. To further evaluate the models’ performance, we introduced the Content Consistency Score (CCS). This metric evaluates the consistency between the model’s outputs and the article content extracted by researchers. The CCS is calculated as follows:

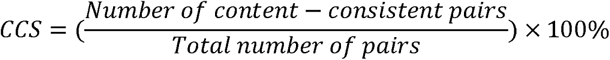

## 3 Results

### 3.1 Description of the information retrieval workflow

The workflow of the proposed RAPID, illustrated in **Figure 1**, is structured into two main components: Input Preparation and Agent Framework. In the Input Preparation phase, two essential preprocessing steps are conducted: Text Preprocessing (detailed in **Methods 2.4**) and Medical Reporting Guidelines Preprocessing. Following this, the agent framework’s core functionality is powered by several GPT-based LLMs for efficient paragraph selection. This framework performs two primary tasks: (1) using GPT-based LLMs to identify paragraphs relevant to a given item and its detailed descriptions, and (2) evaluating the relevance and suitability of selected paragraphs for addressing the items. To improve both efficiency and transparency for the entire system, we employed a series of engineered prompts and structured processes based on previous methodologies ^7,24^. Key aspects of this design include:

1. **Structured prompts:** Prompts are designed to be concise and clear, breaking down each task into well-defined steps. This minimizes the risk of unnecessary or redundant content influencing the LLM’s output.
2. **Item decomposition (ID):** Checklist items are broken into simpler sub-queries, ensuring that the LLMs understand tasks accurately. This method improves both precision and transparency, allowing users to trace each analysis step ^23^.
3. **Multi-round Iterative Paragraph Retrieval Strategy:** Paragraphs are processed in consecutive batches, where each batch builds upon the previous one. This strategy reduces redundancy and improves the precision of paragraph selection ^14^.
4. **Output restriction:** We implemented output constraints, such as JSON format ‘Yes’/‘No’ responses and adjusted output temperature, to improve consistency and facilitate automated processing.

**Figure 1.**
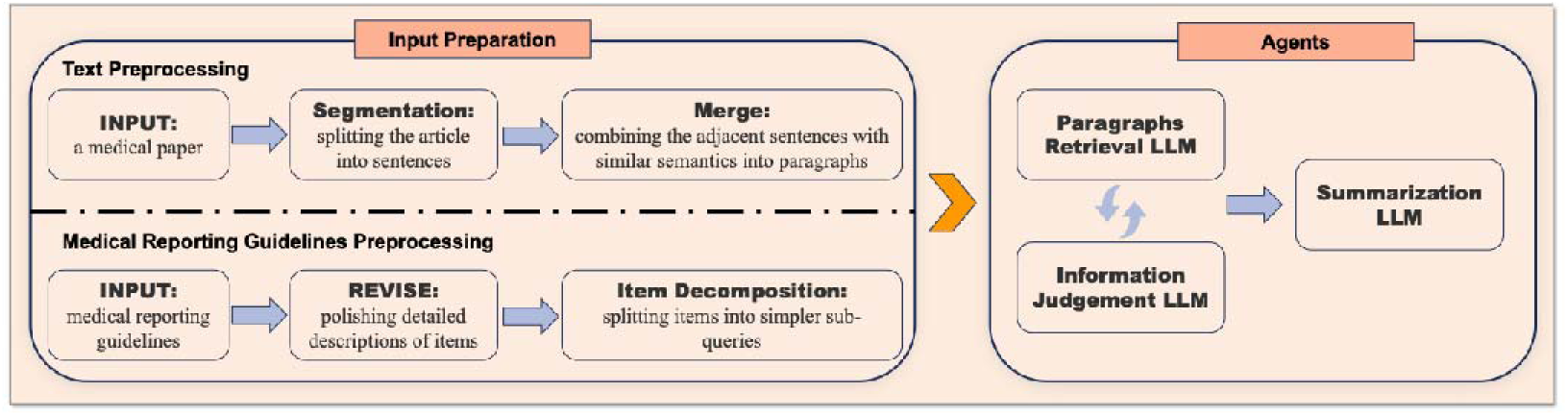
A simplified workflow of RAPID, an automatic information retrieval framework using large language models. This workflow only includes some key components for each step and the fully detailed workflow is shown in Figure 2 and Figure 3.

The detailed steps for polishing item descriptions are depicted in **Figure 2** (A). Item descriptions aim to provide comprehensive explanations of medical report guideline items, facilitating reader understanding by detailing their background and significance. Incorporating well-crafted item descriptions into prompts is beneficial for the subsequent paragraph screening task. However, original descriptions in medical reporting guidelines often contain extraneous or vague content and may lack concrete examples. To address this, we employed LLMs to assess these descriptions against nine evaluation indicators, such as indirectness, readability, and focus (**Supplementary eTable 3** and **eFigure 5**). Descriptions that fell short were iteratively refined based on LLM feedback, and a welly-revised item description will finally be output, usually meeting the set standards within three iterations.

**Figure 2.**
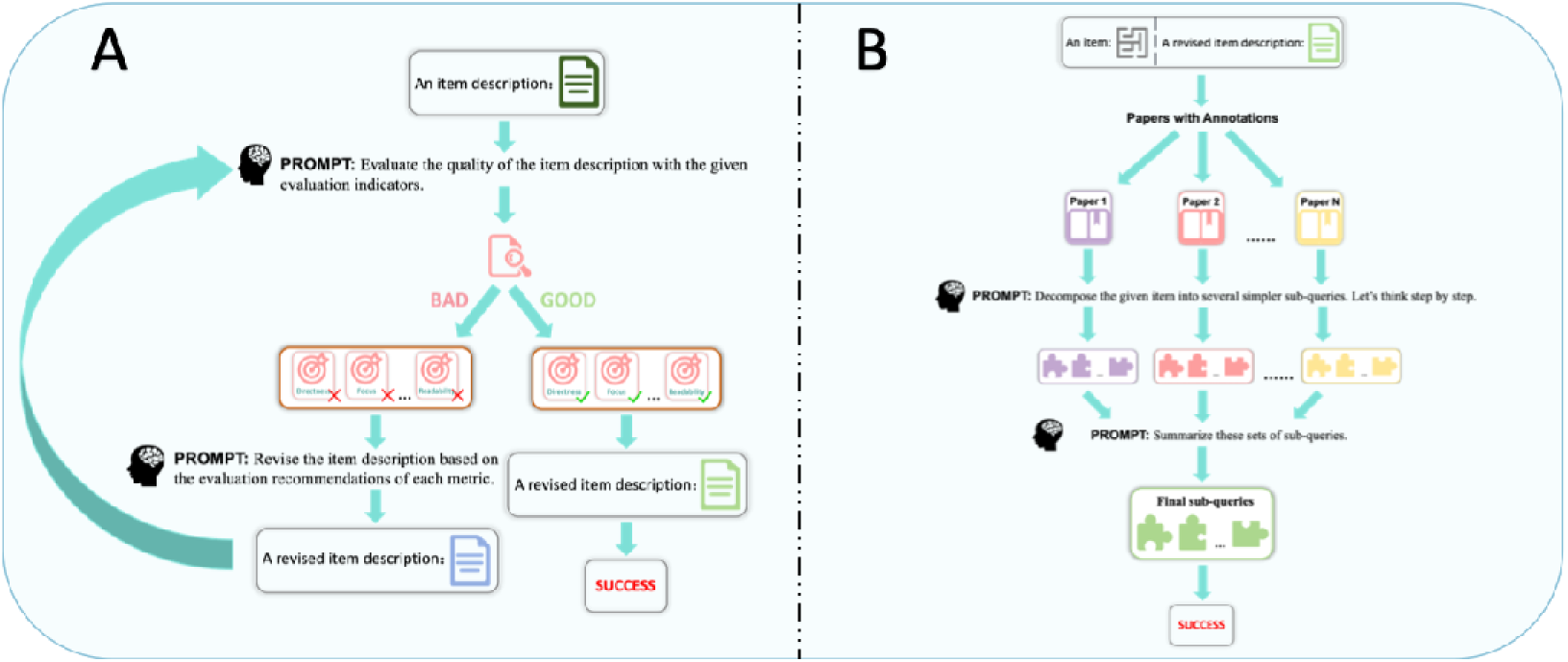
A detailed workflow of RAPID, an automatic information retrieval framework using large language models. (A) Iterative optimization workflow for item description. (B) Workflow for item decomposition.

Once item descriptions were revised, items were decomposed into comprehensible sub-queries (**Figure 2** (B)). Given the diversity of writing styles across articles, we initially employed zero-shot prompting (**Methods 2.2**) to generate multiple sub-query sets for various articles, requiring each sub-query to be related to the previous one while still being answerable independently (**Supplementary eFigure 6**). These sets were then summarized as a final series of sub-queries using summary prompts (**Supplementary eFigure 7**). During experiments, we observed that paper paragraphs corresponding to these sub-queries typically appear in specific sections: early paragraphs aligned with initial sub-queries, while later paragraphs addressed final sub-queries.

Following sub-queries generation, the workflow proceeded to the paragraph screening (**Figure 3** (A)). Rather than directly identifying paragraphs related to the given checklist item, RAPID focused on locating those related to the sub-queries. This approach is straightforward because sub-queries are derived from the original item, so the information retrieved for answering the sub-queries naturally addresses the item itself. To tackle the challenge of identifying relevant content within lengthy articles (the "Needle in a Haystack" problem ^8,12^, paragraphs are grouped into consecutive blocks of 25, with a 5-paragraph overlap to maintain context. The first LLM prompt selects paragraphs within each block, while the second prompt evaluates these selected paragraphs to determine if they could answer the sub-query. If the answer is “No”, the process continues with the next block until an adequate response is found or all blocks are processed. Two key strategies in this process include:

1. Paragraphs selected from previous blocks are included in the next block to avoid unnecessary choices.
2. If no suitable paragraphs are found, additional prompts are used to summarize the information of the intermediate steps to enhance transparency during the final answer generation stage.

**Figure 3.**
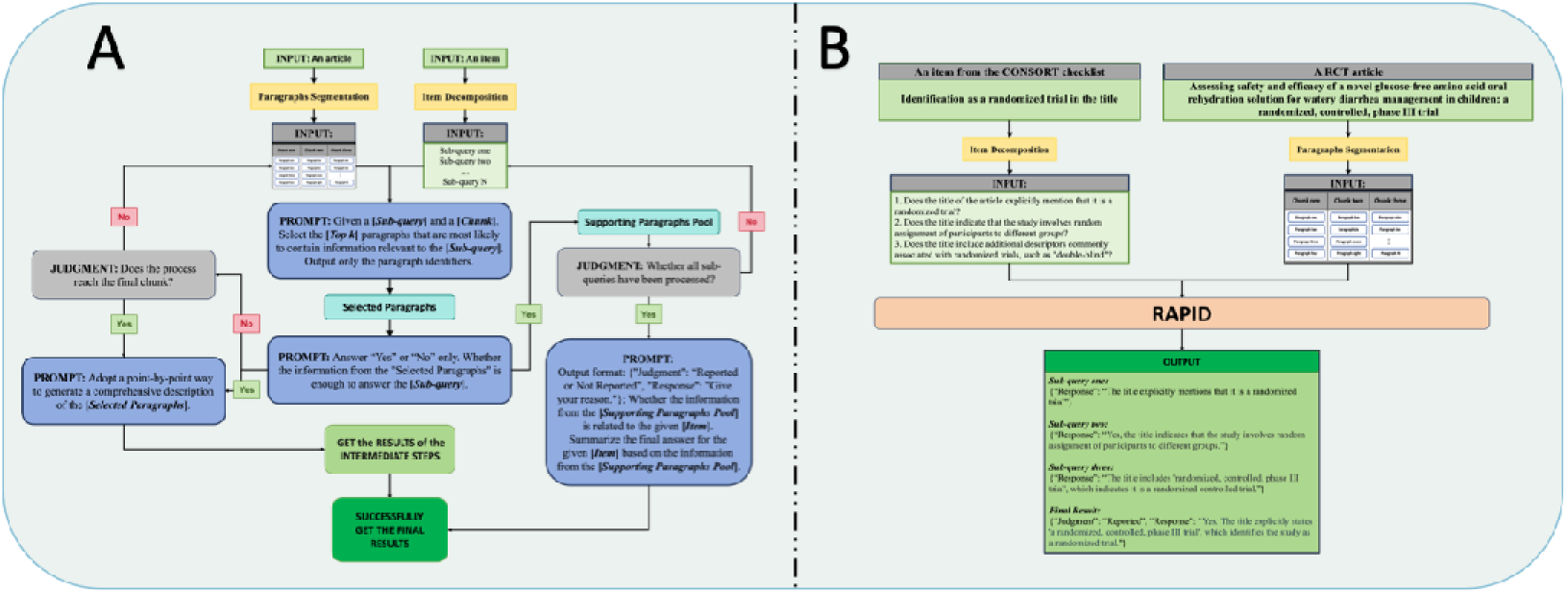
A detailed workflow of RAPID, an automatic information retrieval framework using large language models. (A) LLM agent workflow for paragraph selection and information summary. (B) An example of applying RAPID.

Finally, a structured response prompt (**Supplementary eFigure 8**) generates an organized output (**Figure 3** (B)) based on the retrieved ‘Supporting Paragraphs’. To address potential instruction ambiguity in long-context scenarios, we developed a correction prompt (**Supplementary eFigure 8**) and employed varied temperature settings, ultimately selecting the most frequent response among outputs as the final result.

In our experimentation, we compared RAPID with the Vanilla Prompt method (**Methods 2.3**) using Accuracy and Content Consistency Score (CCS) as evaluation metrics as well as their 95% Confidence Intervals (CI) (**Methods 2.5)**. The LLMs evaluated included ChatGPT-4o, ChatGPT-3.5, Claude-3.5-Sonnet, and deepseek-chat ^1,6,20^. During the experiments, we initially assessed the models performance on the CONSORT dataset, followed by the CONSORT-AI dataset to estimate the scalability of RAPID without any modifications to its core processes. Each model was accessed via its official API using Python 3.10 on a high-performance server (AMD EPYC 7513, Oracle Linux 8.8). For consistent results and reproducibility, we used the following specific model versions: the gpt-4o-2024-05-13 snapshot for ChatGPT-4o, the gpt-3.5-turbo-0125 snapshot for ChatGPT-3.5, the claude-3.5-sonnet-20240620 snapshot for Claude-3.5-Sonnet, and DeepSeek-V2 snapshot for deepseek-chat. To minimize variability in responses, the temperature parameter was set to 0 for all models except deepseek-chat, which, according to its official guidelines, was set to a temperature of 0.7 for tasks requiring nuanced information extraction and analysis. All other parameters were kept at their default values.

### 3.2 Model comparison in assessing compliance of RCTs with CONSORT

#### 3.2.1 Accuracy comparison between different models

To evaluate the effectiveness of our proposed architecture, we calculated accuracy and compared it against the Vanilla Prompt method using various GPT-based LLMs on the CONSORT dataset (**Methods 2.1**). The average accuracy of different methods was summarized in **Table 1**. Experimental results showed that, among the Vanilla Prompt methods, ChatGPT-4o, OpenAI’s most advanced model, achieved the highest accuracy of 84.59% (95%CI = [80.89, 88.60]), significantly outperforming ChatGPT-3.5, which scored 71.62% (95%CI = [65.60, 77.64]). In addition, two other top-performing models, deepseek-chat from Deepseek and Claude-3.5-Sonnet from Anthropic, attained accuracies of around 83.73% (95%CI = [79.63, 87.83]) and 79.89% (95%CI = [74.89, 84.89]), respectively.

**Table 1.** The average accuracy and content consistency score of different models on the CONSORT dataset.

| Models | LLMs | Average Accuracy (%) |  |  | Content Consistency Score (%) |  |  |
| --- | --- | --- | --- | --- | --- | --- | --- |
|  |  | ACC | 95%CI-lower | 95%CI-upper | CCS | 95%CI-lower | 95%CI-upper |
| Vanilla Prompt | ChatGPT-4o<br>(gpt-4o-2024-05-13) | 84.59 | 80.89 | 88.60 | 77.51 | 73.35 | 81.68 |
| Vanilla Prompt | ChatGPT-3.5<br>(gpt-3.5-turbo-0125) | 71.62 | 65.60 | 77.64 | 51.57 | 46.06 | 57.07 |
| Vanilla Prompt | Claude-3.5-Sonnet<br>(claude-3.5-sonnet-20240620) | 79.89 | 74.89 | 84.89 | 69.62 | 64.13 | 75.11 |
| Vanilla Prompt | deepseek-chat<br>(DeepSeek-V2) | 83.73 | 79.63 | 87.83 | 74.27 | 68.99 | 79.55 |
| RAPID | ChatGPT-3.5<br>(gpt-3.5-turbo-0125) | 85.08 | 80.64 | 89.52 | 71.41 | 66.01 | 76.80 |
| RAPID | deepseek-chat<br>(DeepSeek-V2) | 92.11 | 90.0 | 94.22 | 81.14 | 77.15 | 85.12 |

RAPID demonstrated further competitive performance when combined with different models, showcasing its adaptability across various LLMs. When combined with ChatGPT-3.5, RAPID achieved an accuracy comparable to the Vanilla Prompt approach with ChatGPT-4, even exceeding the results of Claude-3.5. Additionally, we extended our evaluation to an open-source model to assess RAPID’s performance. Specifically, we evaluated it with deepseek-chat which ranks higher than ChatGPT-3.5 but slightly below Claude-3.5-Sonnet and ChatGPT-4 in comprehensive assessments ^4^. Despite this ranking, RAPID employed with deepseek-chat achieved an accuracy of 92.11% (95%CI = [90.0, 94.22]), significantly outperforming all previously assessed models and demonstrating its robustness and extensibility. These results indicated that even LLMs with lower performance can match or surpass top-tier closed-source LLMs when integrated with RAPID, highlighting the effectiveness of our method and addressing concerns regarding the use of closed-source models in private system deployments, especially in regions with restrictions. Moreover, as the overall capability of the LLM improved, RAPID’s performance scaled proportionally, indicating its potential for further enhancement. However, due to token budget limitations, we could not fully evaluate the performance of RAPID employed with ChatGPT-4o. Nevertheless, based on the performance trends observed, we are confident that RAPID combined with ChatGPT-4o would deliver superior results.

#### 3.2.2 Consistency of content generated by models compared to humans

The initial evaluation of the models’ performance focused on assessing the consistency of the ‘Reported’/’Not Reported’ classifications with those made by human researchers. In real-world scenarios, different human researchers conducting checklist evaluations may consistently judge certain items as "Reported," but the specific passages they refer to may vary. Therefore, accuracy alone may introduce bias, as it can not reflect how consistently the content produced by models aligns with information extracted by human researchers. To achieve a more comprehensive evaluation of the models’ quality, we further calculated the content consistency score (CCS) (**Methods 2.5**).

As indicated in **Table 1**, the results demonstrated an interesting observation: models can distinguish between ‘Reported’ and ‘Not Reported’ items, yet the basis for their judgment may differ from that of humans. For instance, when using the Vanilla Prompt method with ChatGPT-3.5, a relatively good accuracy of 71.62% is achieved, but CCS is lower at 51.57%. This observation emphasized the significance of CCS as a complementary measure to accuracy.

In addition, this analysis illustrated that RAPID, when combined with deepseek-chat, achieved the highest average CCS at 81.14% (95%CI = [77.15, 85.12]), significantly outperforming other top models. For comparison, the Vanilla Prompt methods achieved lower CCS scores of 77.51% (95%CI = [73.35, 81.68]), 69.62% (95%CI = [64.13, 75.11]), and 74.27% (95%CI = [68.99, 76.8]), when combined with ChatGPT-4, Claude-3.5-Sonnet, and deepseek-chat, respectively. These findings once again exhibited RAPID’s superior performance and its ability to maintain alignment with human judgments, highlighting RAPID as a reliable tool for content extraction tasks, especially in applications that demand consistent and human-aligned outputs.

### 3.3 Extensibility verification of RAPID on CONSORT-AI

To verify the generalizability of RAPID, previously validated on the CONSORT dataset, we applied it to the CONSORT-AI dataset (**Methods 2.1**) without modifying its architecture or prompts. Concurrently, we also evaluated the Vanilla Prompt methods using prompts adjusted for compatibility with the new dataset. All results are presented in **Table 2** and **Supplementary**.

**Table 2.** The average accuracy and content consistency score of different models on the CONSORT-AI dataset.

| Models | LLMs | Average Accuracy (%) |  |  | Content Consistency Score (%) |  |  |
| --- | --- | --- | --- | --- | --- | --- | --- |
|  |  | ACC | 95%CI-lower | 95%CI-upper | CCS | 95%CI-lower | 95%CI-upper |
| Vanilla Prompt | ChatGPT-4o<br>(gpt-4o-2024-05-13) | 73.84 | 60.45 | 87.22 | 62.75 | 49.77 | 75.73 |
| Vanilla Prompt | ChatGPT-3.5<br>(gpt-3.5-turbo-0125) | 61.20 | 49.94 | 72.45 | 36.81 | 28.90 | 44.71 |
| Vanilla Prompt | Claude-3.5-Sonnet<br>(claude-3.5-sonnet-20240620) | 79.82 | 70.60 | 89.04 | 65.19 | 53.61 | 76.77 |
| Vanilla Prompt | deepseek-chat<br>(DeepSeek-V2) | 76.27 | 61.64 | 90.91 | 62.53 | 49.31 | 75.74 |
| RAPID | ChatGPT-3.5<br>(gpt-3.5-turbo-0125) | 76.94 | 69.95 | 83.93 | 64.08 | 54.28 | 73.88 |
| RAPID | deepseek-chat<br>(DeepSeek-V2) | 83.81 | 76.51 | 91.12 | 72.51 | 62.67 | 82.34 |
| RAPID<br>(w/o) | deepseek-chat<br>(DeepSeek-V2) | 81.60 | 73.77 | 89.43 | 71.84 | 61.96 | 81.72 |

Within the Vanilla Prompt methods, ChatGPT-4o just achieved an accuracy of 73.84% (95%CI = [60.45, 87.22]) and a CCS of 62.75% (95%CI = [49.77, 75.73]), significantly lower than that of Claude-3.5-Sonnet (ACC = 79.82%, 95%CI = [70.60, 89.04]; CCS = 70.29%, 95%CI = [53.61, 76.77]). This result contrasted with prior findings on the CONSORT dataset, where ChatGPT-4o outperformed Claude-3.5-Sonnet, highlighting considerable variability in the Vanilla Prompt method’s performance when models and prompts are adjusted. This observation suggested that the Vanilla Prompt method’s effectiveness is sensitive to model and dataset changes, indicating a degree of instability and unpredictability across different configurations.

For RAPID, when combined with ChatGPT-3.5, it achieved an accuracy of 76.94% (95%CI = [69.95, 83.93]) and a CCS of 64.08 (95%CI = [54.28, 73.88]), comparable to that of Vanilla Prompt method with closed-source LLMs. Additionally, when combined with deepseek-chat, a more powerful LLM, RAPID demonstrated a significantly better performance (ACC=83.81%, 95%CI = [76.51, 91.12]; CCS=72.51%, 95%CI = [62.67, 82.34]), which align with the conclusions from the previous case study and effectively verify RAPID’s robustness and potential for broader application to other medical reporting guidelines.

### 3.4 Web application display

RAPID was deployed on an online application accessible at http://rapid.advancedguides.ai, offering four main functions: (1) Document Upload; (2) Feedback Generation for Medical Reporting Guidelines; (3) Explanation of Feedback; and (4) Automated Report Generation. These functions are embedded within an intuitive user interface, as illustrated by a sample screenshot of an RCT study in **Supplementary eFigure 9**. This figure showcases two primary interactive pages within the application: the Document Upload page and the Article Display page.

On the Document Upload page (**Supplementary eFigure 9** (A)), users have access to the following capabilities: (1) Personal Online Storage Access: Users can view and manage their personal document storage directly within the platform; (2) Document Upload: Users can upload documents in PDF or Word format and specify a medical reporting guideline; (3) Article Navigation: Users can select a processing completed articles from their database and navigate to the Article Display page for detailed interaction.

The Article Display page, depicted in **Supplementary eFigure 9** (B), provides an interactive environment for reviewing and refining report content. Here, users can: (1) View Checklist Items: A checklist panel on the right side of the page presents items that are color-coded for quick reference: green indicates reported items, while red denotes items that are missing. Users can click on each item’s number to highlight the corresponding paragraphs in the document for easier identification. (2) Access Explanation of System Judgments: By clicking any item on the checklist, users can view a rationale, which includes sub-queries related to the system’s judgment. (3) Make Corrections: If users believe the system’s feedback requires adjustment, users can correct the judgment by clicking the correction button located in the bottom right corner of each item. (4) Automated Report Generation: By clicking the download button at the top right of the page, the system can automatically generate the checklist report, integrating both system and user-provided insights.

## 4 Discussion

The rapid expansion of medical literature and the ongoing evolution of reporting guidelines^11^ highlighted the growing need for automated tools to assess compliance with these checklists. Existing tools, such as CONSORT-NLP^27^, used NLP technology and reported up to 90% accuracy. However, this tool was limited, covering only 34 of the 37 CONSORT checklist items, providing no explanatory support for its judgments, and requiring specific training. In contrast, our proposed method encompassed all items, offered detailed explanations for each ‘Reported’/’Not Reported’ judgment, and was capable of adapting to various reporting guidelines without requiring specific training.

In our experiments, we observed that nearly all methods scored poorly on two items such as items 2 and 4 of the CONSORT-AI checklist (**Supplementary eFigure 3)**. Detailed analysis of these results revealed two primary phenomena behind these low scores: (1) For item 2, researchers tended to mark ‘Reported’ when patient screening procedures were mentioned in the RCTs. In contrast, LLMs were more stringent, only marking ‘Reported’ when specific details about the inclusion and exclusion criteria of the input data were present; and (2) For item 4, LLMs consistently marked ‘Reported’ when they detected any information about the source of the AI intervention method. However, researchers only considered this ‘Reported’ if a clear version number was provided. In addition, we analyzed instances where our method provided incorrect answers and discovered a pattern: while our architecture successfully retrieved paragraphs containing relevant information, the model made errors in the final summarization. We believe this is due to the limited summarization capabilities of the LLMs. We are confident that as large language models continue to improve, this issue will be effectively resolved in the future.

To alleviate this situation currently, we introduced a secondary correction mechanism which involves integrating common LLMs’ errors (**Supplementary eFigure 8**) into the prompt for adjustment and using varied temperature settings. Ultimately, we select the most frequent response among the outputs as the final result. Although our method has achieved over 80% accuracy on two real-world datasets, we recommend users perform a secondary correction when using it, as our method is primarily intended to expedite the process. Last but not least, our experimentation revealed that the Vanilla Prompt method is highly sensitive to configuration changes, leading to inconsistent outputs. In contrast, RAPID maintained stable performance due to four key features: (1) efficient compression of contextual length, (2) a sequential paragraph selection process guided by previously selected paragraphs, (3) carefully designed prompts with reasoning steps, and (4) the secondary correction mechanism.

To support researchers in evaluating medical reports, we developed a web-based application based on our methods. This application provided reminders for complex checklist items by displaying the reasoning path and allowed users to modify the system’s final results if needed (**Results 3.4** for details). Our application offered distinct advantages for multiple user groups: (1) Medical Authors: Enhance report quality by verifying manuscript completeness and automatically generating submission reports, simplifying the submission process; (2) Medical Researchers: Facilitate evaluation of report quality within specific fields, identify research gaps, and help shape future studies; (3) Medical Editors: Verify the accuracy and relevance of submission checklists, expediting manuscript distribution; (4) Reviewers: Assist in manuscript assessment by identifying reporting deficiencies and providing deeper insights for review comments. RAPID addresses the growing need for automated reporting compliance tools and supports the advancement of medical reporting guidelines. By promoting integrity, transparency, and interpretability in clinical trials, RAPID holds promise for substantial contributions to the medical field.

## 5 Conclusion

In this study, we introduced RAPID, an automated information extraction framework designed to improve the efficiency of evaluating medical reports across various reporting guidelines. To support adaptability and scalability evaluation, we developed two datasets, respectively, for CONSORT and CONSORT-AI checklists. RAPID demonstrated state-of-the-art performance on both datasets. Specifically, the system achieved an average accuracy of 92.11% and a content consistency score of 81.14% on the CONSORT dataset, and an accuracy of 83.81% with a content consistency score of 72.51% on the CONSORT-AI dataset. The success of RAPAD lies in a sequence of well-engineered prompts and carefully crafted structured processes, precisely identifying relevant content while eliminating redundant content from extensive text. Furthermore, to support researchers, we developed a web-based application built upon this RAPID, providing a user-friendly interface for streamlined access to RAPID’s capabilities.

### Limitation

We acknowledge that our work has some limitations. Firstly, the prompts used in this study were specifically optimized for ChatGPT, which may not be ideal for other LLMs. Additionally, the architecture had relatively high token consumption, suggesting room for optimization in this area. Furthermore, we observed performance variability across different LLMs when summarizing the final answer for certain items, even when provided with the correct reference content. This problem to some extent can be mitigated through secondary correction, as demonstrated by our approach, or by combining the results of multiple models. It will be fully addressed in the future by improving the capabilities of LLMs. In the future, we would like to continuously optimize the overall architecture, reduce token usage, and extend our research to cover more medical reporting guidelines.

## 6 Acknowledgments

This article was partly revised by ChatGPT-4o, a large language model from OpenAI. The model was only used to improve the readability and language quality of the article. The version of ChatGPT-4o used was gpt-40-2024-05-13. The exact prompt used with the model was "[Sentence] Please help me revise the sentence". It was used in several parts of the article, including but not limited to the introduction and methods. It was used under strict human supervision and control. Additionally, authors carefully reviewed and polished the content generated by the model. The model was only used to improve the language and readability of the article and was not used to copy or replace any researcher tasks, such as generating scientific insights, analyzing and interpreting data, or drawing scientific conclusions.

