## Supplementary for "RAPID: Reliable and efficient Automatic generation of submission rePorting checklists with Large language moDels"

### Content

- **eTable 1. CONSORT unique checklist.**
- **eTable 2. CONSORT-AI unique checklist.**
- **eTable 3. Nine metrics for revising item description.**
- **eFigure 1. Accuracies of different methods for various items on the CONSORT dataset.**
- **eFigure 2. Content consistency scores of different methods for various items on the CONSORT dataset.**
- **eFigure 3. Accuracies of different methods for various items on the CONSORT-AI dataset.**
- **eFigure 4. Content consistency scores of different methods for various items on the CONSORT-AI dataset.**
- **eFigure 5. Prompt for revising item description.**
- **eFigure 6. Prompt for item decomposition.**
- **eFigure 7. Prompt for summarizing sub-queries.**
- **eFigure 8. Prompts for multi-agent framework.**
- **eFigure 9. A Randomized Controlled Trial case study - interactive interface for streamlined research document compliance evaluation.**

### eTable 1. CONSORT unique checklist.

| No. | Item |
|---|---|
| 1 | Identification as a randomized trial in the title. |
| 2 | Structured summary of trial design, methods, results, and conclusions (for specific guidance, see CONSORT for abstracts). |
| 3 | Scientific background and explanation of rationale. |
| 4 | Specific objectives or hypotheses. |
| 5 | Description of trial design (such as parallel, factorial) including allocation ratio. |
| 6 | Important changes to methods after trial commencement (such as eligibility criteria), with reasons. |
| 7 | Eligibility criteria for participants. |
| 8 | Settings and locations where the data were collected. |
| 9 | The interventions for each group with sufficient details to allow replication, including how and when they were actually administered. |
| 10 | Completely defined pre-specified primary and secondary outcome measures, including how and when they were assessed. |
| 11 | Any changes to trial outcomes after the trial commenced, with reasons. |
| 12 | How sample size was determined. |
| 13 | When applicable, explanation of any interim analyses and stopping guidelines. |
| 14 | Method used to generate the random allocation sequence. |
| 15 | Type of randomization; details of any restriction (such as blocking and block size). |
| 16 | Mechanism used to implement the random allocation sequence (such as sequentially numbered containers), describing any steps taken to conceal the sequence until interventions were assigned. |
| 17 | Who generated the random allocation sequence, who enrolled participants, and who assigned participants to interventions. |
| 18 | If done, who was blinded after assignment to interventions (for example, participants, care providers, those assessing outcomes) and how. |
| 19 | If relevant, description of the similarity of interventions. |
| 20 | Statistical methods used to compare groups for primary and secondary outcomes. |
| 21 | Methods for additional analyses, such as subgroup analyses and adjusted analyses. |
| 22 | For each group, the number of participants who were randomly assigned received intended treatment, and were analyzed for the primary outcome. |
| 23 | For each group, losses and exclusions after randomization, together with reasons. |
| 24 | Dates defining the periods of recruitment and follow-up. |
| 25 | Why the trial ended or was stopped. |
| 26 | A table showing baseline demographic and clinical characteristics for each group. |
| 27 | For each group, number of participants (denominator) included in each analysis and whether the analysis was by original assigned groups. |
| 28 | For each primary and secondary outcome, results for each group, and the estimated effect size and its precision (such as 95% confidence interval). |
| 29 | For binary outcomes, presentation of both absolute and relative effect sizes is recommended. |

| No. | Item |
|---|---|
| 30 | Results of any other analyses performed, including subgroup analyses and adjusted analyses, distinguishing pre-specified from exploratory. |
| 31 | All important harms or unintended effects in each group (for specific guidance, see CONSORT for harms). |
| 32 | Trial limitations, addressing sources of potential bias, imprecision, and, if relevant, multiplicity of analyses. |
| 33 | Generalizability (external validity, applicability) of the trial findings. |
| 34 | Interpretation is consistent with results, balancing benefits and harms, and considering other relevant evidence. |
| 35 | Registration number and name of trial registry. |
| 36 | Where the full trial protocol can be accessed, if available. |
| 37 | Sources of funding and other support (such as the supply of drugs), role of funders. |

### eTable 2. CONSORT-AI unique checklist.

| No. | Item |
|---|---|
| 1 | Explain the intended use of the AI intervention in the context of the clinical pathway, including its purpose and its intended users (for example, healthcare professionals, patients, public). |
| 2 | State the inclusion and exclusion criteria at the level of the input data. |
| 3 | Describe how the AI intervention was integrated into the trial setting, including any onsite or offsite requirements. |
| 4 | State which version of the AI algorithm was used. |
| 5 | Describe how the input data were acquired and selected for the AI intervention. |
| 6 | Describe how poor quality or unavailable input data were assessed and handled. |
| 7 | Specify whether there was human–AI interaction in the handling of the input data, and what level of expertise was required of users. |
| 8 | Specify the output of the AI intervention. |
| 9 | Explain how the AI intervention's outputs contributed to decision- making or other elements of clinical practice. |
| 10 | Describe results of any analysis of performance errors and how errors were identified, where applicable. If such analysis was planned or done, justify why. |
| 11 | State whether and how the AI intervention and/or its code can be accessed, including any restrictions to access or re-use. |

### eTable 3. Nine metrics for revising item description.

| Evaluation Metrics | Description |
|---|---|
| Enhanced Clarity and Conciseness | Description should eliminate unnecessary details that don't contribute to the primary message. For instance, specific citations and references within the body of the description are removed if they don't add significant value to the reader's understanding. This makes the content |

| | |
|---|---|
| | more accessible, especially to readers who may not be familiar with the detailed background literature. |
| **Evaluation Metrics** | **Description** |
| Focused Content | Description should directly address the question or item at hand without deviating into tangential details. This focused approach ensures that the essential information is immediately clear to the reader, facilitating better comprehension of the core concepts. By directly stating the most important points without embedding them in dense paragraphs, ensure that these key messages are not overlooked by the reader . |
| Improved Readability | Description should avoid breaking words across lines and use a more structured layout. This results in improved readability, making it easier for readers to follow the narrative without getting lost in complex sentence structures or unnecessary jargon. |
| Absence of Redundant Information | Redundant explanations and justifications that do not directly contribute to answering the question or clarifying the item should be removed. This streamlining of content helps in maintaining the reader's interest and ensures that the text remains engaging. |
| User-Centric Explanation | Description should be tailored to be more user-centric, focusing on how the information presented will affect or be useful to the reader, especially in terms of understanding trial procedures, outcomes, and the practical implications. |
| Transparency and Replicability | Description should emphasize the importance of transparency and replicability in trial procedures, making it a point to mention when and how information can assist in these endeavors. This is done by providing clear and detailed instructions or descriptions that can be easily followed or understood by others in the field. |
| Use of Specific Examples | Descriptions should include specific, real-world examples to illustrate the application and impact of the AI intervention, enhancing understanding through practical scenarios. |
| Clinical Focus | The narrative should emphasize the clinical relevance and implications of the AI intervention, highlighting its significance for patient care and decision-making processes. |
| Acknowledgment of Limitations and Risks | The style should openly address the potential errors, limitations, and risk mitigation strategies associated with the AI intervention, promoting ethical reporting and trust. |

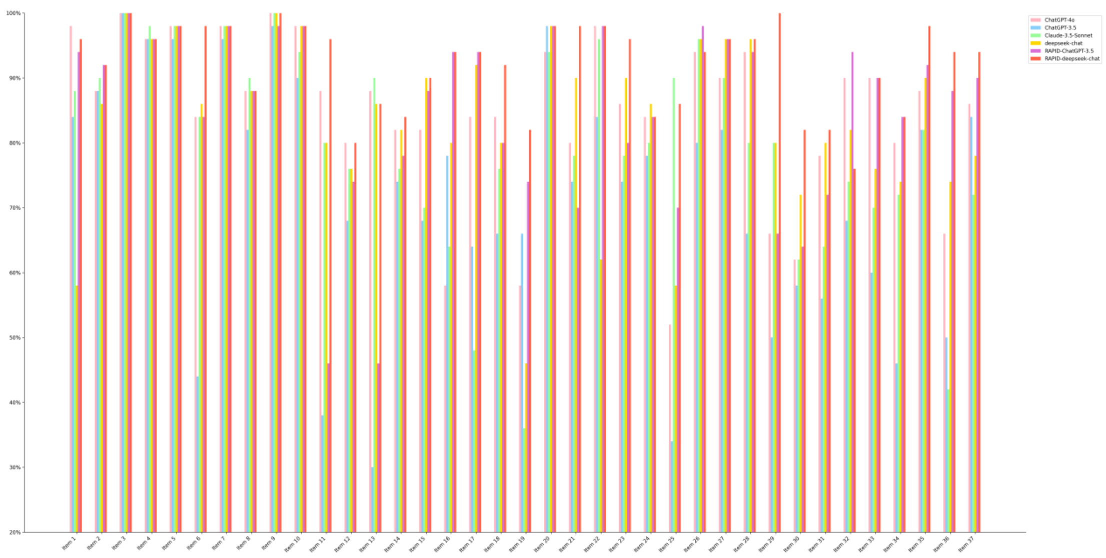


**eFigure1. Accuracies of different methods for various items on the CONSORT dataset.**

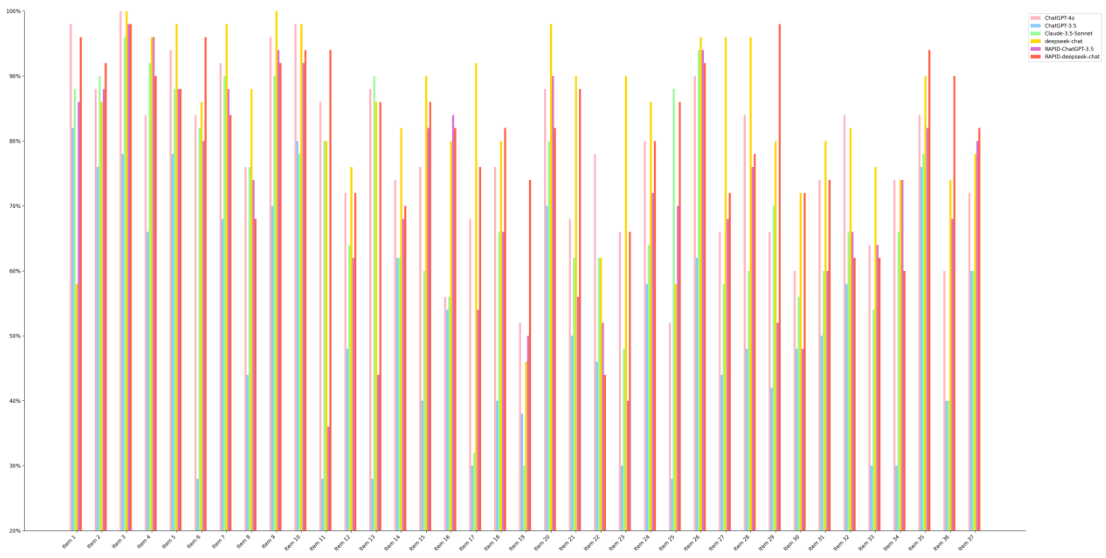


**eFigure2. Content consistency scores of different methods for various items on the CONSORT dataset.**

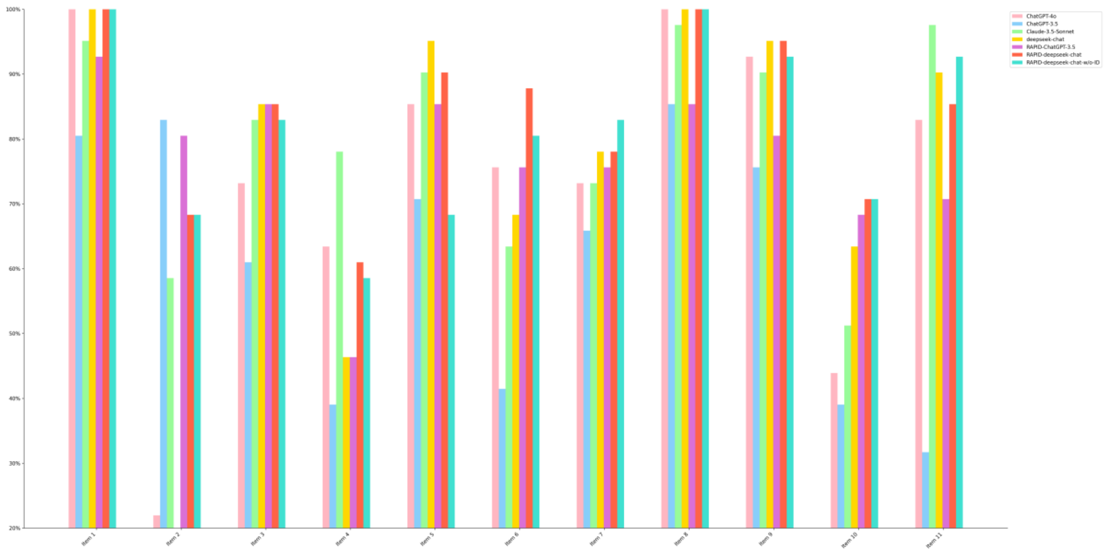


**eFigure3. Accuracies of different methods for various items on the CONSORT-AI dataset.**

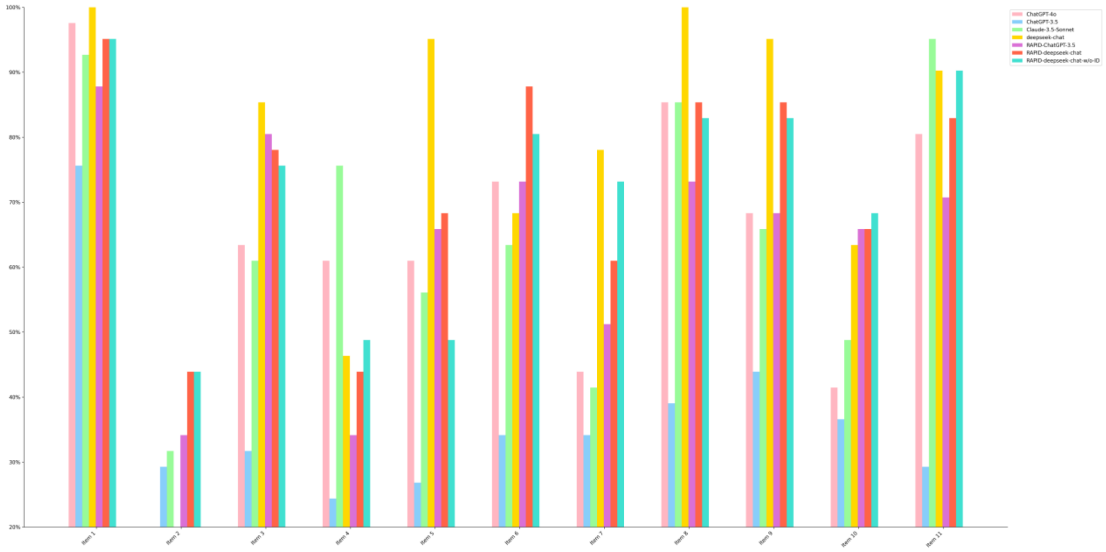


**eFigure4. Content consistency scores of different methods for various items on the CONSORT-AI dataset.**

Prompt for revising item description (Part A)

````
### Your role:
```start
Suppose you are a clinical trial author and evaluator.
```end

### Evaluation metrics:
```start
{evaluation metrics}
```end

### Your task:
```start
I will provide you an item and specify the reporting guidelines checklist it originates from, a question for the given item, and a detailed description
for the question.
Your task is to evaluate the descriptive style of the detailed description based on each metric outlined in the 'Evaluation Metrics' section, with the
scoring for each metric ranging from 1 to 10 points.
```end

### Item:
```start
{item}
```end

### Question:
```start
{question}
```end

### Detailed description:
```start
{description}
```end


### Response template:
```start
1. "Enhanced Clarity and Conciseness": [Score]
2. "Focused Content": [Score]
3. "Improved Readability": [Score]
4. "Absence of Redundant Information": [Score]
5. "User-Centric Explanation": [Score]
6. "Transparency and Replicability": [Score]
7. "Use of Specific Examples": [Score]
8. "Clinical Focus": [Score]
9. "Acknowledgment of Limitations and Risks": [Score]
```end

### Detailed description evaluation scores (response must start with ```start and end with ```end, follows the "Response template"):
````

Prompt for revising item description (Part B)

````
### Your role:
```start
Suppose you are a clinical trial author and evaluator.
```end

### Evaluation metrics:
```start
{evaluation metrics}
```end

### Item:
```start
{item}
```end

### Question:
```start
{question}
```end

### Detailed description:
```start
{description}
```end

### Detailed description evaluation scores:
```start
{evaluation_scores}
```end

### Reminder:
```start
1. The score of each metric listed in "Evaluation metrics" ranges from 0 to 10, with 10 being the highest.
2. "Detailed description evaluation scores" provide a metric-based assessment and targeted recommendations for enhancing the "Detailed Description."
```end

### Your task:
```start
Your task is to revise the "Detailed Description" according to the "Evaluation Metrics" and the improvement suggestions within the "Detailed
Description Evaluation Scores." Your goal is to produce an optimized detailed description that would achieve higher scores across all metrics.
```end


It is your turn to perform the task. Ensure that the "Optimized detailed description" is less than 300 words and does not include a summary.
### Optimized detailed description (response must start with ```start and end with ```end):
````

**eFigure5. Prompt for revising item description.**

**Prompt for item decomposition**

````
### Uploaded article:
<document>
{uploaded_article}
</ducoment>

### Reminder:
```start
1. The content of the "Uploaded article" is framed by <document> and </document>.
2. The "Question text" provides the question that needs to be answered.
3. The "Question description" provides additional and detailed explanations for the "Question text".
4. The "Question answer" provides the answer to the question from the "Uploaded article". If it contains None, that means the answer does not exist in
the "Uploaded article".
```end

### Your role:
```start
Suppose you are a clinical author and evaluator.
```end

### Your task:
```start
Your task is to effectively decompose complex, multi-hop questions into simpler, manageable sub-questions or tasks. This process involves breaking down
a question that requires information from multiple sources or steps into smaller, more direct questions that can be answered individually.
Here's how you should approach this:
Analyze the "Question text": Carefully read the multi-hop question to understand its different components. Identify what specific pieces of information
are needed to answer the main question.
```end

Format your answer as multiple lines of text. Ensure that each subsequent question follows from the previous one is self-contained and is capable of
being answered on its own. Ensure there is exactly one line break between each line and each line must start with "```start" and end with "```end".

Now it's your turn to answer:
### Question text:
```start
{question_text}
```end
### Question description:
```start
{question_description}
```end
### Question answer (original text in paper):
```start
{question_answer}
```end
### Decomposed quires (Let's think step by step):
````

**eFigure6. Prompt for item decomposition.**

**Prompt for summarizing sub-queries**

````
### Reminder:
```start
1. The "Question text" provides the question that needs to be answered.
2. The "Question description" provides additional and detailed explanations for the "Question text".
3. The "Decomposed quires" consist of simpler, more manageable decomposed questions for the complex "Question text" from {num} different articles.
"Decomposed quires" is logically structured, each subsequent question follows from the previous one is self-contained, and is capable of being answered
on its own.
```end

### Your role:
```start
Suppose you are a clinical trial author and evaluator.
```end

### Your task:
```start
"Decomposed quires" consist of simpler, more manageable decomposed questions for the complex "Question text" from {num} different articles. "Decomposed
quires" is logically structured, each subsequent question follows from the previous one is self-contained, and is capable of being answered on its own.
Your task is to refine and summarize the "Decomposed queries" from the {num} examples into a coherent and logically structured set of queries. This
process involves making a concise and coherent summary of the "Decomposed queries", the summary contains more logical, smaller, direct questions that
can be answered individually.
Here's how you should approach this:
1. Analyze the "Decomposed quires" and carefully read the "Question text" and "Question description".
2. Identifying how to answer sub-questions step by step can ultimately get the answer to the "Question text".
3. Summarize the "Decomposed quires" into a coherent and logically structured set of queries. Each subsequent question follows from the previous one is
self-contained and is capable of being answered on its own. The "Summarized decomposed quires" maximally generates 6 decomposed queries.
```end

Format your answer as multiple lines of text.
Ensure that each subsequent question follows from the previous one is self-contained and is capable of being answered on its own. Ensure that each sub-
question does not contain any content which is only related to the corresponding example. Ensure there is exactly one line break between each line and
each line must start with "```start" and end with "```end".

Now it's your turn to answer:
### Question text:
```start
{question_text}
```end
### Question description:
```start
{question_description}
```end
### Decomposed quires:
```start
{decomposed_quires}
```end
### Summarized decomposed quires: (Let's think step by step):
````

**eFigure7. Prompt for summarizing sub-queries.**

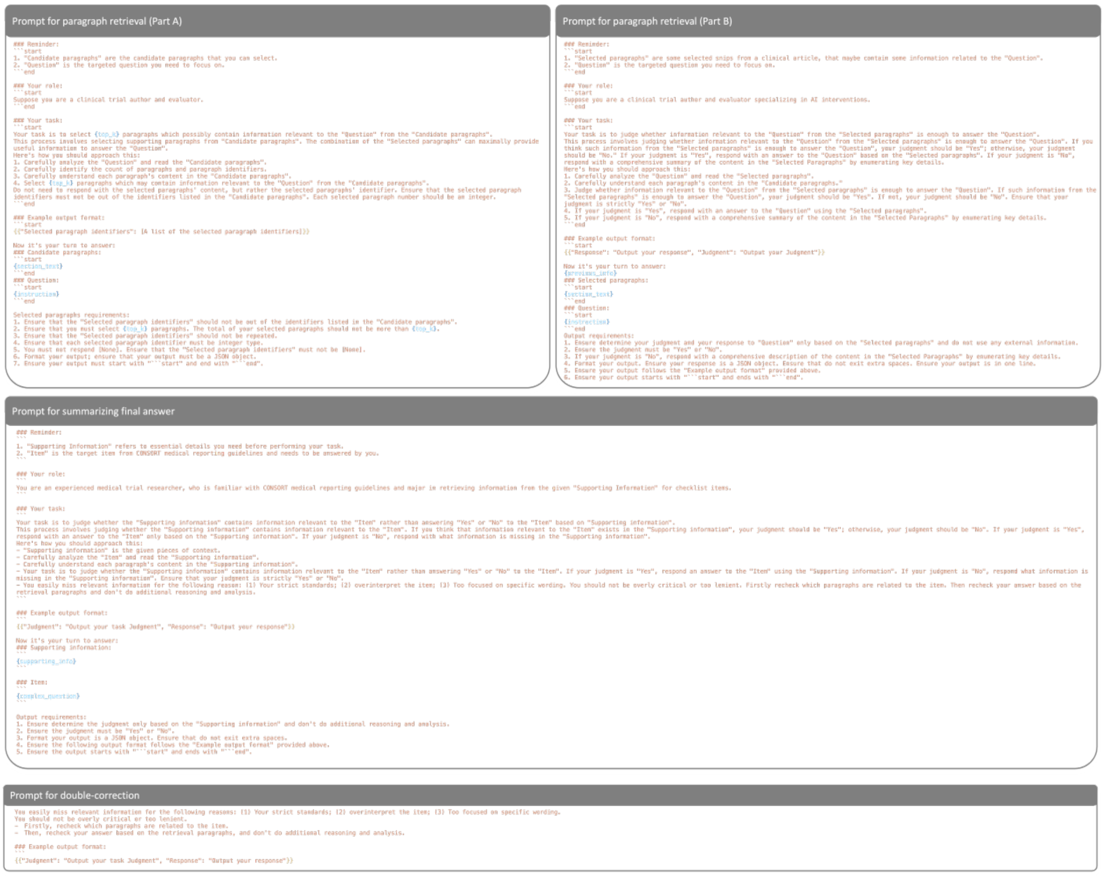


**eFigure8. Prompts for multi-agent framework.**

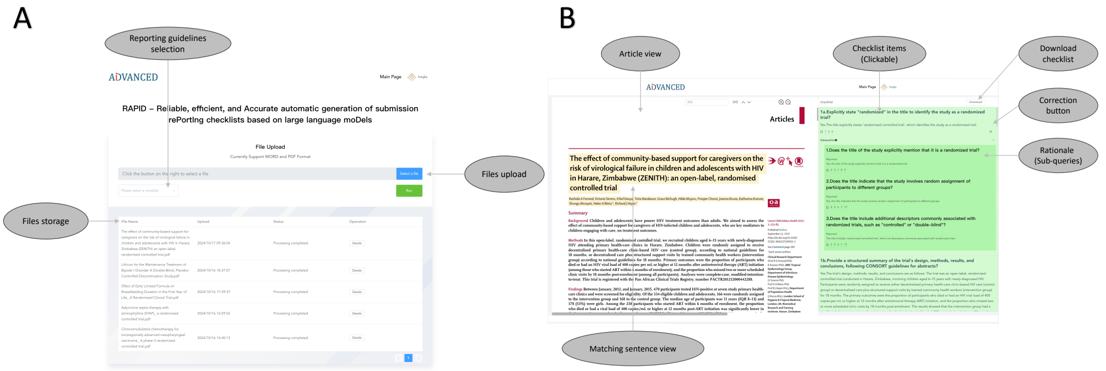


**eFigure9. A Randomized Controlled Trial case study - interactive interface for streamlined research document compliance evaluation. Screenshot A shows the Document Upload page with options for file upload, file storage, and reporting guideline selection. Screenshot B illustrates the Article Display page with four main panels: (1) Article view; (2) Matching sentence view; (3) Checklist items; and (4) Rationale**
